# Efficient Detection of Communities in Biological Bipartite Networks

**DOI:** 10.1101/105197

**Authors:** Paola Pesántez-Cabrera, Ananth Kalyanaraman

**Author notes:** The authors are with the School of Electrical Engineering and Computer Science, Washington State University, Pullman, WA, 99164.

**Keywords:** Heterogeneous biological data, bipartite networks, graph algorithms, community detection, bipartite modularity

## Abstract

Methods to efficiently uncover and extract community structures are required in a number of biological applications where networked data and their interactions can be modeled as graphs, and observing tightly-knit groups of vertices (“communities”) can offer insights into the structural and functional building blocks of the underlying network. Classical applications of community detection have largely focused on unipartite networks—i.e., graphs built out of a single type of objects. However, due to increased availability of biological data from various sources, there is now an increasing need for handling heterogeneous networks which are built out of multiple types of objects. In this paper, we address the problem of identifying communities from biological *bipartite networks*—i.e., networks where interactions are observed between *two different types* of objects (e.g., genes and diseases, drugs and protein complexes, plants and pollinators, hosts and pathogens). Toward detecting communities in such bipartite networks, we make the following contributions: i) (*metric*) we propose a variant of bipartite modularity; ii) (*algorithms*) we present an efficient algorithm called *biLouvain* that implements a set of heuristics toward fast and precise community detection in bipartite networks; and iii) (*experiments*) we present a thorough experimental evaluation of our algorithm including comparison to other state-of-the-art methods to identify communities in bipartite networks. Experimental results show that our *b*iLouvain algorithm identifies communities that have a comparable or better quality (as measured by bipartite modularity) than existing methods, while significantly reducing the time-to-solution between one and four orders of magnitude.

## 1 Introduction

THE increasing identification and characterization of genes, protein complexes, diseases, and drugs have highlighted a need to incorporate *heterogeneity* while analyzing complex biological data. A heterogeneous network is composed of multiple types of objects. Identifying “community” structures that transcend data boundaries in such a network could provide new insights that may not be readily visible by examining only a specific data type in isolation. For instance, identifying a group of genes that have been implicated across a set of diseases could possibly reveal hidden links among seemingly different diseases or disease conditions, and in the process help identify new drugs and therapies [1]. Similarly, identifying active gene clusters across different subsets of brain regions could provide new insights into brain function [2].

Graph-theoretic representations offer a natural way to model networks built out of heterogeneous data. This work focuses on *bipartite networks* as a way to model the interplay between two different data types. Bipartite networks are those which have two types of vertices such that edges exist only between vertices of the two different types. Bipartite networks are called *unweighted* when the edges between vertices are either present or not. Alternatively, if edges are given different values representing the strength of the associations among vertices the bipartite network is called *weighted*. Some examples of biological bipartite networks include (but are not limited to) gene-disease [3], gene-drug [4], plants-pollinators [5], and host-pathogen [6].

While the idea of modeling heterogeneity as a graph problem is not necessarily new *per se*, algorithm development efforts have been more recent [7], [8]. Once modeled as a bipartite network, we can view the problem of identifying cluster structures between the two different data types as a problem of *community detection in bipartite networks*. For example, in the case of the gene-drug network we could identify groups of drugs that might inhibit or otherwise modulate groups of genes. We also could detect groups of genes that are suitable for drug repurposing but may not currently have a drug targeting them. Similarly, for plantpollinator networks we could identify groups of pollinators that limit or promote the establishment and persistence of plant species.

Given a graph, the goal of community detection is to partition the set of vertices into “communities” such that vertices that are assigned to the same community have a higher density of edges among them than to the vertices in the rest of the network. Community detection can be used to reveal hidden substructures within real world networks, without any known prior knowledge on either the number or sizes of the output communities.

Community detection is a well studied problem in literature [9]. However, the treatment of the problem on bipartite networks has been sparse. Because edges connect vertices of two different types, the classical definition of communities [10] does *not* directly apply. Projection-based approaches typically result in loss of the bipartite structural information [11]. Instead, the notion of communities needs to be redefined so that a community of vertices of one type is formed on the basis of the strength of its shared connections to vertices of the other type (as shown in Fig. 1).

**Fig. 1.**
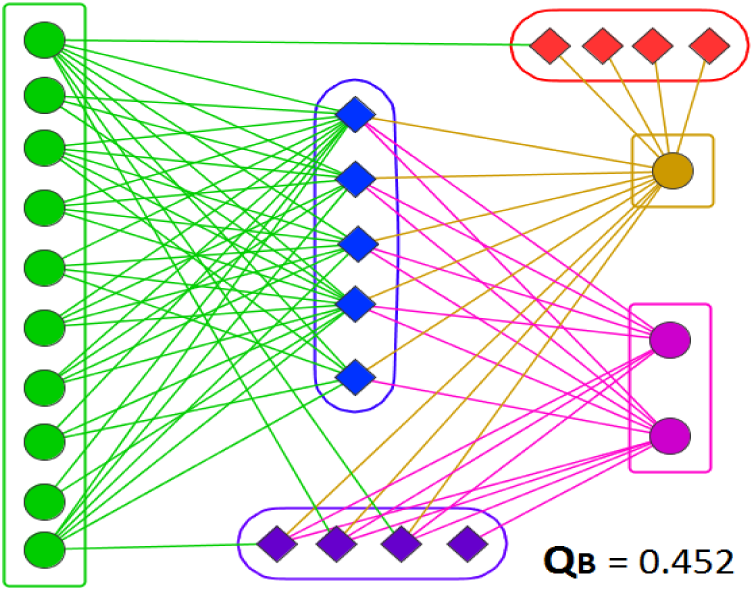
Communities detected by our *biLouvain* algorithm in the *bezerra2009* bipartite network [5] of 13 Malpighiaceae oil-flowers (circles) vs. 13 oil-collecting bees (diamonds). Nodes of the same color correspond to the same community. *Q*_*B*_ stands for bipartite modularity.

To unveil important new associations, evaluation of the goodness of a community-wise division of a bipartite network also becomes critical. To this end, a measure called *bipartite modularity* can be defined by extending the classical measure for unipartite networks [10]. Modularity is a statistical measure which calculates the difference between the observed fraction of intra-community edges to an expected fraction in an equivalent random graph — i.e., null model. A higher value of modularity suggests a clearer community structure within the underlying network. Modularity optimization is an NP-Hard problem [12]; however, a number of efficient heuristics are used in practice [9]. For bipartite networks, multiple formulations of bipartite modularity have been proposed [13], [14], [15], [16]. Of these, Murata’s definition is known to overcome some of the limitations of the other definitions (see Sections 2 and 4).

### 1.1 Contributions

We present a direct, efficient method (*biLouvain*) to optimize our modified version of Murata’s modularity (*Murata+*). Specifically, the main contributions of this paper are:

- ***Murata+ modularity:*** We initially consider the classical definition of Murata’s modularity [15] and discuss its suitability for community detection in Section 4. In the process of evaluation, we identify an inconsistency in the classical formula and subsequently propose a simple variant, which we call *Murata+*.
- ***biLouvain algorithm:*** We present efficient algorithmic heuristics for optimized detection of bipartite communities using the *Murata+* modularity (Section 5). Our approach extends the Louvain algorithm [17], which is one of the most efficient and widely used heuristics for unipartite networks. Consequently, we call our algorithm *biLouvain*. As part of our algorithm, we provide ways to calculate the modularity gain resulting from vertex migrations in a bipartite network — a step that constitutes the core of the Louvain heuristic.
- ***Experimental evaluation:*** We present a thorough experimental evaluation of our algorithm using both synthetic and real world networks and in comparison to other state-of-the-art methods (Section 6). Experimental results show that our algorithm identifies communities that have a comparable or better quality (as measured by the Murata+ bipartite modularity) than existing methods, while significantly reducing the timeto-solution between one and four orders of magnitude.

A preliminary version of the work presented on this paper has been published in [18]. The rest of this paper is organized as follows. Section 2 describes the key related work for community detection in the context of bipartite networks. Section 3 presents the basic notation and terminology for use throughout the paper. Section 4 explores the applicability and extension of Murata’s modularity as a way to determine the quality of a particular division of a network into communities. Section 4 describes in detail the algorithm we have developed—i.e., *biLouvain* algorithm, for detection of communities in bipartite networks. Section 6 presents the experimental evaluation of our algorithm on different real and synthetic networks. Section 7 concludes the paper.

## 2 Related Work

### Modularity

Community detection has been extensively studied in the context of unipartite graphs [9]. Most of these algorithms use variants of the modularity measure, as defined by Newman [10]. Multiple efforts have extended the classical definition of modularity to bipartite networks:

i) Guimerà *et al.* [14] defines bipartite modularity as the cumulative deviation from the random expectation of the number of edges between vertex members of the same bipartite community. The main weakness of this definition is that it focuses on connectivity from the perspective of only one vertex type.
ii) Barber [13] extends the scope to include connectivity information from both vertex types. However, this definition has a limitation of enforcing a one-to-one correspondence between the communities from both vertex types — i.e., the number of communities should be equal on both sides.
iii) Murata’s definition [15] overcomes the above limitations by not enforcing a one-to-one mapping between the communities of either side; more details of this measure are presented in Section 4.1.
iv) Suzuki-Wakita [16] uses a “prominence factor” that averages the volume of connections to all communities.

While there is no general consensus on the modularity definition to use for bipartite networks, we chose to adopt the Murata’s definition (with modifications as described in Section 4.2). Experimental results (Section 6) demonstrate the high quality of results produced by this definition.

### Bipartite community detection

As for community detection in bipartite networks, there have been a handful of efforts so far. In 2006, Newman [21] proposed a special case of spectral algorithm called *Leading Eigenvector* for unipartite networks; however it has been used for bipartite networks as well through a conversion of the bipartite matrix into its unipartite representation. Here the Laplacian matrix is replaced with a modularity matrix that is shown to have the same properties. This method divides the network in two communities by choosing the eigenvector with the largest eigenvalue and the process continues until the leading eigenvalue is zero or negative.

In 2007, Barber [13] developed two modularity-based algorithms, viz. *BRIM* and *Adaptive BRIM*. The latter uses the BRIM algorithm; however the number of communities, *c*, gets doubled at each iteration until there is no further gain in modularity. If the current value of *c* causes modularity to decrease, the heuristic performs a bisection search for *c* in the interval wherein the maximum modularity lies.

In 2009, Liu and Murata [19] proposed a hybrid algorithm called *LPBRIM* that uses the Label Propagation heuristic to search for a best community configuration, and subsequently uses BRIM to refine the results. This approach was extended and improved in 2016 by Beckett [20] in a tool called *DIRTLPAwb+*.

In 2014, Larremore *et al.* [22] developed an approach (*biSBM*) that uses a Stochastic Block model to maximize a likelihood function. This approach assumes that the number of output communities should be known *a priori*.

Table 1 summarizes the conceptual differences among these methods, relative to our algorithm *biLouvain* that is proposed in this paper.

**TABLE 1.**
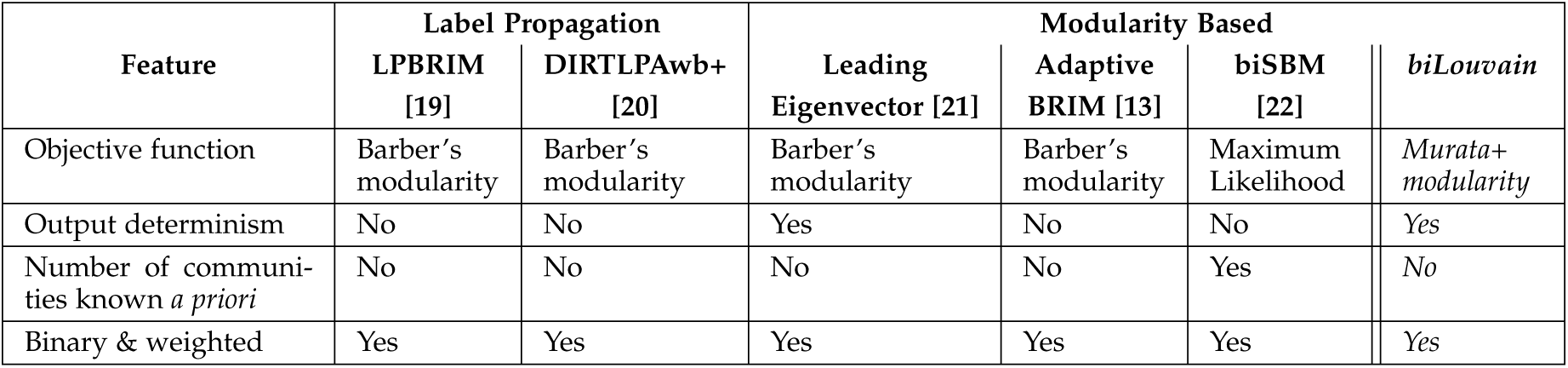
A comparison of bipartite community detection methods.

## 3 Basic Notation and Terminology

We will use *G*(*V*_1_ ∪ *V*_2_, *E*, *ω*) to denote an undirected bipartite graph. Here, *V*_1_ and *V*_2_ represent the two sets of vertices, and *E* represents the set of edges such that each edge *e* = (*i, j*) ∈ *E* is such that *i* ∈ *V*_1_ and *j* ∈ *V*_2_. Each edge *e* is also associated with a numerical weight *ω*(*e*). We will assume that the edge weights are non-negative values that reflect the strength of the relation between any two vertices. The sum of the weights of all edges incident on a vertex *i* is said to be its *weighted degree* (denoted by γ(*i*)). Additionally, we use the term *binary networks* to describe networks whose edges are unweighted. In such cases, all edges that exist are assumed to have a unit weight.

Let *n*_1_ and *n*_2_ denote the number of vertices in *V*_1_ and *V*_2_ respectively, and *m* denote the number of edges. Let *M* = 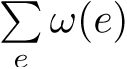. For sake of consistency, we use *i*’s to denote vertices in *V*_1_, and *j*’s for vertices in *V*_2_.

A *community* represents a subset of either *V*_1_ or *V*_2_. For ease of exposition, we use *C*’s to denote communities taken from *V*_1_ and *D*’s to denote communities taken from *V*_2_. Let *P*_1_ = {*C*_1_, *C*_2_, …, *C*_*k*1_} denote a set of communities in *V*_1_ such that it represents a partitioning of *V*_1_. Similarly, let *P*_2_ = {*D*_1_, *D*_2_, …, *D*_*k*_2__} represent a set of communities in *V*_2_ such that it represents a partitioning of *V*_2_. Throughout this paper, we assume *k*_1_ need *not* be equal to *k*_2_.

In our algorithmic discourse, we denote the present community containing any vertex *i* ∈ *V*_1_ as *C*(*i*), and the present community containing any vertex *j* ∈ *V*_2_ as *D*(*j*).

## 4 Bipartite Modularity

### 4.1 Murata’s Bipartite Modularity

In what follows, we describe Murata’s modularity [15] although using our own notation for convenience.

Given a pair of communities, *C* ∈ *P*_1_ and *D* ∈ *P*_2_, and let *e* denote any edge that connects a vertex in community *C* with a vertex in community *D*. Consequently, we define:

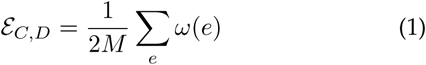

Note that by this definition, ɛ_*C,D*_ = ɛ_*D,C*_. Also note that the term 2*M* corresponds to the sum of the weighted degree of all vertices^1^ — i.e., 2*M* = ∑*_i_ γ(i)*. We use the term *A*_*C*_ to denote the fraction of this term contributed by a given community *C*.

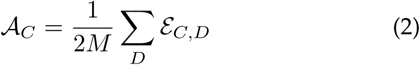

Furthermore, we define the *co-cluster mate* of a community *C* to be a community *D* ∈ *P*_2_ to which *C* has the most concentration of its edges — i.e.,

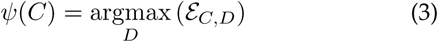

We refer to the ordered tuple 〈*C*, *ψ(C)*〉 as a *co-cluster*. Fig. 2 illustrates this concept.

**Fig. 2.**
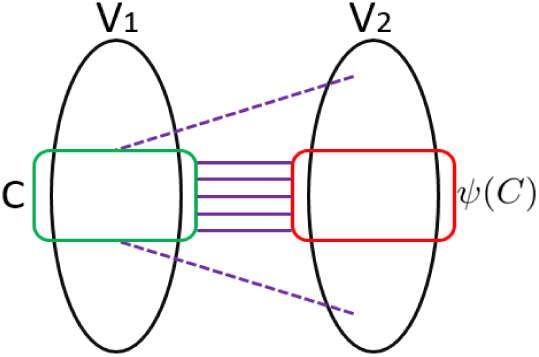
Illustration of a co-cluster 〈*C*, *ψ(C)*〉 where *ψ(C)* is refered as the co-cluster mate of community *C*.

Similarly, the *co-cluster mate* of a community *D* is defined as follows:

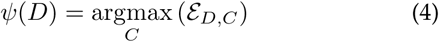

The ordered tuple 〈*D*, *ψ(C)*〉 is also a *co-cluster*.

#### Definition 1.

Given a bipartite graph *G*(*V*_1_ ∪ *V*_2_, *E*, ω), and two sets of communities *P*_1_ in *V*_1_ and *P*_2_ in *V*_2_, Murata’s bipartite modularity *Q*_*B*_ is defined as follows [15]:

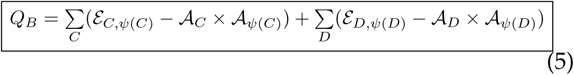

Intuitively, Murata’s modularity is calculated by pairing every community from one side with a community on the other side that it has maximum connections to. The first term inside the two summations in Eqn. 5 corresponds to the fraction of such “intra-cocluster” edges. The second term inside each of the summations is the expected fraction of such edges in a randomly generated bipartite graph with an identical vertex degree sequence. As in Newman’s modularity [10] for unipartite networks, the idea is to encourage a partitioning that maximizes intra-cocluster edges while discouraging a partitioning that groups unrelated vertices.

### 4.2 *Murata+*: Proposed Bipartite Modularity

From Eqns. 3, 4, and 5 we make the following two observations:

**Observation 1.** If a community *C* picks a community *D* as its co-cluster mate (by Eqn. 3), *D* need *not* necessarily pick *C* (by Eqn. 4).
**Observation 2.** The statistical terms *𝒜*_*C*_ × *𝒜*_*ψ*(*C*)_ and *𝒜*_*D*_ × *𝒜*_*ψ*(*D*)_ are used in the final modularity calculation of Eqn. 5, but they are *not* used while picking the co-cluster mates (Eqns. 3 and 4).

Observation 1 implies that the co-cluster relationship is *nonsymmetric*. This relaxation is necessary to avoid a method-enforced one-to-one mapping between communities and their co-cluster mates. However, the relaxation could also lead to an undesirable effect of a potential lack of cohesion between communities and their co-cluster mates. For bipartite networks, we typically attempt to explain the grouping of a community based on its co-cluster. While one-to-one mapping would make it too restrictive for this purpose, it is also important *not* to make it excessively many-to-many. For most practical inputs, a middle ground is more desirable where the expected mapping remains closer to an one-to-one mapping. For instance, we can expect a strong (if not strict) two-way correlation between a set of genes and the set of diseases they impact.

Observation 2 indicates a matter of inconsistency because a community *C* picks a co-cluster mate solely based on the positive term, while the final modularity is calculated taking into account also the negative term. At best, this can lead to an overestimated value for modularity *Q*_*B*_, when a co-cluster mate is selected. More importantly, we argue that the negative term is in fact essential as otherwise it could potentially lead to a scenario where a community and its co-cluster mate could be of vastly different sizes. This is shown in Fig. 3.

**Fig. 3.**
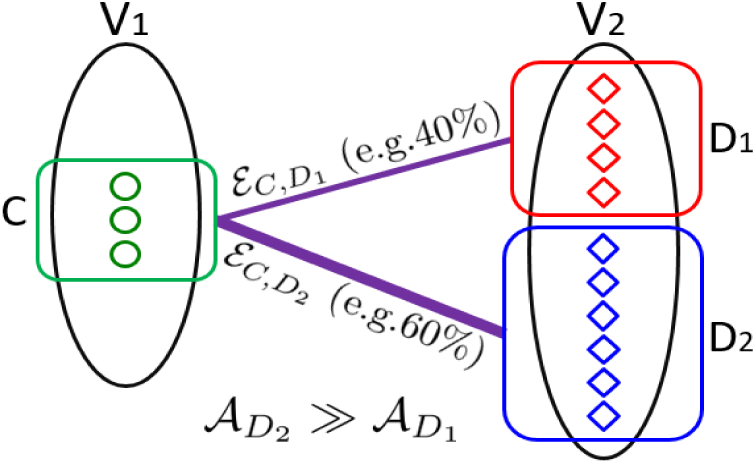
An illustrative example of a case where a community *C* has, say 40% of its edges connected to community *D*_1_ and the remaining 60% of its edges connected to community *D*_2_. Under this scenario, *C* has two choices for its co-cluster mate, **ψ*(C)* = *D*_1_ and **ψ*(C)* = *D*_2_, with *ε*_*C,D*_1__ being only marginally smaller than *ε*_*C,D*_2__ but the sizes of *C* and *D*_2_ could be vastly different (as shown). In scenarios like this, ignoring the negative term may have the undesirable effect of picking a co-cluster mate that is significantly different in size. A better choice of a co-cluster mate for *C* is *D*_1_, not only because of its comparable size, but also because it is likely to lead to a better modularity.

To address the above inconsistency issue within the classical definition, we propose a variant of bipartite modularity by simply redefining the *co-cluster mate* selection criterion to include the negative term:

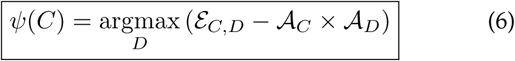

Similarly, a *co-cluster mate* of a community *D* is defined as follows:

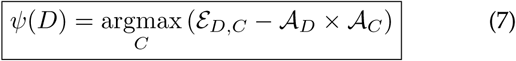

The modularity expression is the same as Eqn. 5. It should be clear that this revised definition would make the choice of co-cluster mates consistent with the modularity calculation — i.e., fixing the problem with Observation 2.

In addition, the revised definition preserves the nonsymmetry property (Observation 1), while being better positioned than the classical definition to encourage an one-toone mapping, wherever possible, without strictly enforcing it. This is because of the reduced degree of freedom that a community is likely to have (with the introduction of the negative term) when selecting its co-cluster mate.

Henceforth, we refer to our revised version of the Murata’s modularity as *Murata+* (Eqns. 5, 6, and 7), and use it as our primary objective function.

## 5 Bilouvain: An Algorithm for Bipartite Community Detection

In this section, we present our *biLouvain* algorithm for community detection in bipartite networks. We adapt the widely used *Louvain* heuristic [17] to work for bipartite networks. While *biLouvain* follows the same algorithmic template provided by Louvain, it differs in the objective function (by using *Murata+*) and in the way all the key steps are computed, as will be elaborated below.

Like the original algorithm, *biLouvain* is a multi-phase, multi-iterative algorithm, where each *phase* is a series of *iterations*, and the algorithm moves from one phase to another using a graph compaction step. Within every iteration, each vertex makes a local decision on its community based on a net modularity gain function. The main steps of the *biLouvain* algorithm are as follows (see Fig. 4):

1) Given an input bipartite graph *G*(*V*_1_ ∪ *V*_2_, *E*, *ω*), initialize a set of *n*_1_ + *n*_2_ communities, where *n*_1_ = *|V*_1_| and *n*_2_ = *|V*_2_|, by placing each vertex in its own community.
2) At every iteration, all vertices in *V*_1_ and *V*_2_ are scanned linearly (in an arbitrary order). For each vertex *i*:
  a) Acquire a list of its **candidate communities** to which it can potentially migrate to;
  b) Evaluate the **modularity gain** that would result from the scenario of migrating *i* to each of the candidate communities.
  c) Finally, migrate vertex *i* to a candidate community that maximizes the modularity gain, only if such gain is positive (otherwise, no change).
3) A phase ends when the net modularity gain achieved between two consecutive iterations is negligible — i.e., below a certain threshold *τ*_*i*_, which we refer to as the *iteration cutoff*.
4) Once a phase terminates, a new graph 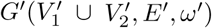 is generated through a **compaction step**, which collapses each community to a vertex, and edges and their weights in the new graph corresponds to the strength of edges connecting any two communities.
5) The new compacted graph is input to the next phase (step 1). The algorithm terminates when any two consecutive phases result in a negligible modularity gain, defined by a threshold *τ*_*p*_, which we call *phase cutoff*.

**Fig. 4.**
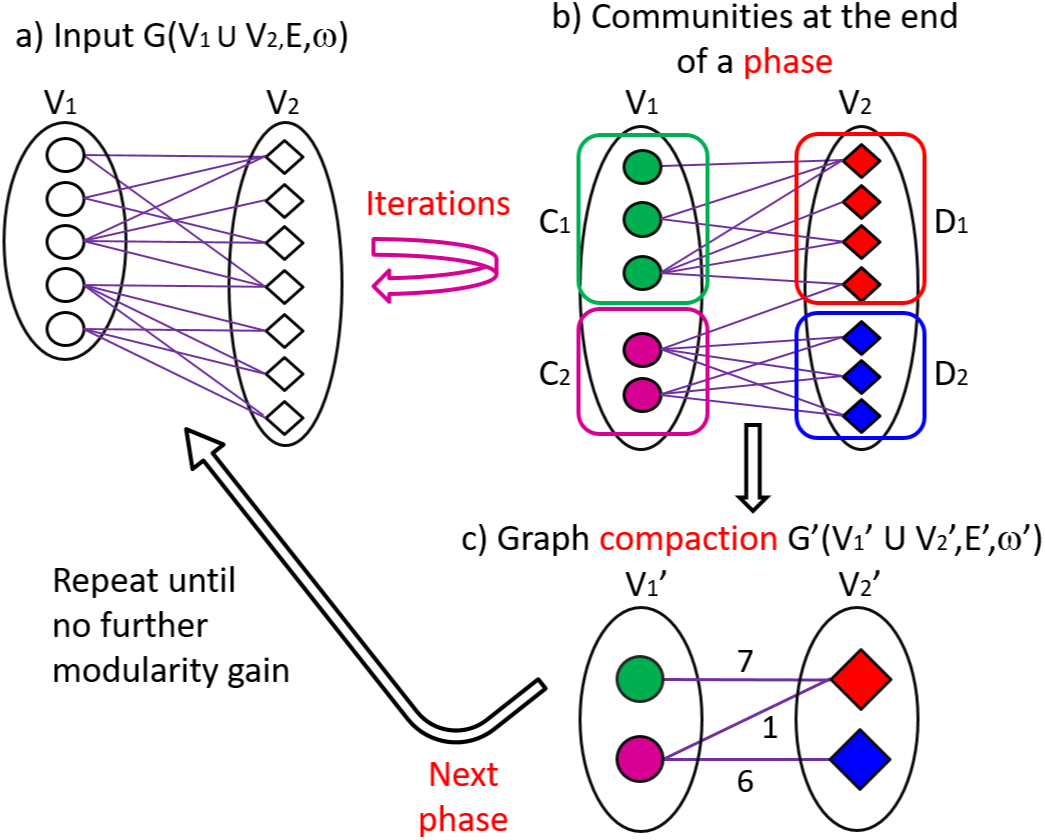
An example to illustrate the main steps of our algorithm for community detection: *biLouvain*. a) The input bipartite graph. Vertices of the two different types are shown in two different shapes. b) After a sequence of iterations, the biLouvain algorithm converges (based on net modularity gain) and a phase is completed. The figure shows the vertices and the communities they belong to at the end of the phase. In this simple example, communities *C*_1_ and *D*_1_ are co-cluster mates to one another, and *C*_2_ and *D*_2_ are co-cluster mates to one another. c) The graph is compacted by collapsing each detected community into a new “vertex” and collapsing inter-community edges between every pair of communities into new “edges” with corresponding weights. This compacted graph is input to the next phase until convergence.

In what follows, we describe how the key steps that are impacted by the bipartite structure are implemented in *biLouvain*. More specifically, the classical definition for candidate communities and their computation (step (2a)), the expression for calculating the modularity gain resulting from a vertex migration, and the algorithm to calculate the modularity gain (step (2b))—all of these need to be defined taking into account the bipartite structure.

### 5.1 Computing Candidate Communities

A *candidate community* of a vertex is a community to which that vertex can potentially migrate at any given iteration of the *biLouvain* algorithm, with a realistic chance of accruing a positive modularity gain.

For ease of exposition, we explain the process of computing candidate communities from the point of view of a vertex *i* in *V*_1_. It should be easy to see that the same approach works for any vertex *j* ∈ *V*_2_.

For a given vertex *i* ∈ *V*_1_, let Γ(*i*) denote the set of neighbors of *i* in *V*_2_ — i.e.,

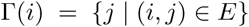

Let Γ’(*i*) denote the set of vertices in *V*_1_ that the neighbors of *i* are connected to.

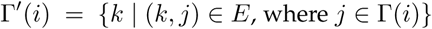

Consequently, the set of candidate communities for *i*, *Cand*(*i*) is given by:

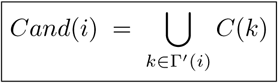

Intuitively, a vertex *i* ∈ *V*_1_ can only migrate to communities in *V*_1_, within which it has at least one 2-hop neighbor (i.e., via its vertex neighbors in *V*_2_). Moving to any other community in *V*_1_ (i.e., not in this candidate set) will result in a decrease in modularity.

### 5.2 Calculating Modularity Gain

In the case of unipartite networks [17], calculating the expected modularity gain resulting from moving a vertex from one community to another can be executed in constant time if appropriate data structures are maintained. In the case of bipartite networks, this is not the same because of the following two lemmas (as shown in Fig. 5):

**Fig. 5.**
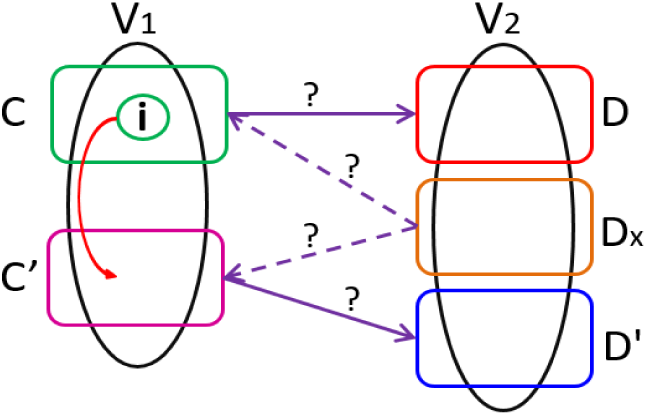
Illustration of vertex *i* evaluating its migration from community *C* to community *C’*. If this move were to be executed, the co-cluster mate choices for both *C* and *C’* (presently, *ψ*(*C*) = *D* and *ψ*(*C’*) = *D’*) could potentially change (Lemma 5.1). Furthermore, the co-cluster mate choice for an arbitrary community *Dx* which has at least one edge incident from either *C* or *C’* could also potentially change (Corollary 5.2). Consequently, a question mark is used to show scenarios where the corresponding co-cluster mate assignments need to be re-evaluated.

#### Lemma 5.1.

If vertex *i* moves from *C* to *C’*, then the choice of co-cluster mates for either community could change.

***Proof:*** The migration of vertex *i* from *C* to *C’* affects the values 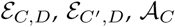 and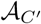, for any *D*, in Eqn. 6. □

Note that the lemma applies to migrations of vertices in *V*_2_ as well.

Define the community set *N* (*C*) as follows:

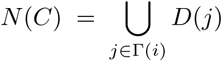

The above lemma leads to the following corollary:

#### Corollary 5.2.

If vertex *i* moves from *C* to *C’*, then the cocluster mate choices for any of the communities in *N* (*C*) or in *N* (*C’*) could potentially change.

***Proof:*** Due to the migration of *i* from *C* to *C’*, any community *D*_*x*_ of *V*_2_ to which either of these two communities is connected (see Fig. 5) could potentially change its cocluster mate preference with changes to the right hand side values of its corresponding *ψ*(*D*_*x*_) calculation (i.e., inside Eqn. 7). This implies that *D_x_ ϵ N* (*C*) *∪ N* (*C’*). □

It should be intuitively clear why these two lemmas should be true. In short, the two equations for co-cluster mate choices (Eqns. 6 and 7) depend on both the positive and the negative terms, and either of those two terms could change for the communities covered in Lemma 5.1 and Corollary 5.2 as a result of *i* moving out of *C* into *C’*. It should also be clear that the co-cluster mate choices for no other communities in the rest of the graph are affected by this move.

Based on the above lemma and corollary, we constitute a set containing all *affected communities* from *i’s* migration from *C* to *C’*—i.e., *S* = {*C, C’}* ∪ *N* (*C*) ∪ *N* (*C’*). Subsequently, we compute the overall net modularity gain for this single vertex move as follows:

We calculate the contribution of a community *D* and its co-cluster mate *ψ(D)* to the summation in the modularity Eqn. 5 as follows:

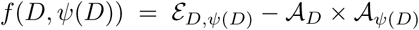

Consequently, the change in a community *D*’s contribution to the overall modularity *Q*_*B*_ due to the change in its co-cluster mate (from *ψ(D)* to *ψ(D)*’ imposed by vertex *i*’s move) is given by:

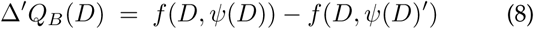

Finally, the overall modularity gain is given by:

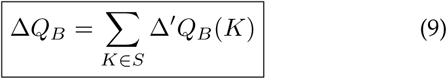

This modularity gain is calculated for each possible vertex move into one of its candidate communities; and finally, vertex *i* is moved to that candidate community which maximizes the gain (assuming it is positive).

### 5.3 Complexity Analysis

The run-time within an iteration is dominated by the time taken to calculate the modularity gain for all the vertices (Section 5.2). In our implementation we keep efficient data structures to enable us to calculate each community’s contribution to the overall modularity, in constant time. Given this, the worst-case run-time complexity for calculating the maximum modularity gain for a given vertex *i* at any given iteration is 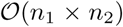—this is for the worst-case scenario of all *C* communities connected to all *D* communities.

As all vertices are linearly scanned within each iteration, the worst-case run-time complexity is 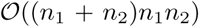 per iteration—which makes our exact algorithm a cubic algorithm. In practice; however, we can expect inputs to be sparse—that would imply a quadratic behavior in the initial iterations; but as the algorithm progresses, the number of communities can only shrink and along with it, also the run-time per iteration.

### 5.4 Performance heuristics

In what follows, we present a collection of heuristics aimed at improving the performance of the *biLouvain* algorithm in practice.

#### 5.4.1 Vertex Ordering

In the *biLouvain* algorithm, the order in which vertices are processed could potentially impact the performance of the algorithm and the quality of community-wise partitioning to varying degrees, depending on the input and on the ordering scheme used.

To understand the impact of vertex ordering on performance, note that the community assignment made for a vertex on one partition (say *V*_1_) at any given iteration is dependent on the community states of its neighboring vertices in the other partition (*V*_2_), and also on the community states in the same partition (for selecting candidate communities). Also note that initially the number of communities on each partition is equal to its number of vertices. When coupled together, these observations imply that if vertex decisions are all made, say sequentially, within one partition prior to the other partition, then the time taken for processing vertices on the first partition is likely to be significantly higher than for the vertices in the second partition during the same iteration. However, this performance impact is expected to diminish in the later iterations of the algorithm as communities get larger, thereby shrinking the number of communities. Consequently, vertex ordering is likely to have an effect on the algorithm’s performance.

As for quality, the impact of vertex ordering is likely to be relatively less. It can be expected that for real world inputs with well-defined community structures, the state of communities typically converges faster in the first few iterations of the algorithm, while largely remain stable in the later iterations. However, the final output quality could still differ based on the vertex ordering used.

To evaluate this quality-time tradeoff imposed by vertex ordering, we implement these vertex ordering schemes:

1) **Sequential:** Within each iteration, the vertices in one partition (say, *V*_1_) are all processed (in some arbitrary order) prior to vertices in the other partition.
2) **Alternate:** Within each iteration, the processing of vertices from the two partitions is interleaved—i.e., alternating between the two partitions, until one of the partitions is exhausted at which point the algorithm defaults to the sequential mode to cover the remaining vertices of the larger partition.
3) **Random:** Within each iteration, the order of processing vertices in *V*_1_ ∪ *V*_2_ is randomized. By fixing the random seed, one can ensure that the output remains deterministic across multiple runs on the same input.

Another contributing factor to dictate an ordering scheme’s impact on performance is the constituent vertex partition sizes. If the two partitions are of skewed sizes (e.g., *n*_1_ ≪ *n*_2_) then the Random scheme is expected to have an advantage in run-time over the other two schemes, whereas if the two partitions have comparable sizes, then all three schemes can be expected to behave similarly.

#### 5.4.2 Structural Properties

The performance of the algorithm can be further improved by observing and taking advantage of certain structural motifs (properties) within bipartite networks and their relation to the modularity expression. Such structural motifs can include simple topological features such as a star (i.e., vertex *i* in *V*_1_ is connected to *k* vertices in *V*_2_), a chain (i.e., a linked list) inside a bipartite network, and common co-cluster mate (i.e., communities *C*_1_, *C*_2_ ∈ *P*_1_ have community *D* ∈ *P*_2_ assigned as their co-cluster mate), or more complex attributes such as a k-core or inexact versions of chains and stars embedded within a larger subgraph.

The idea is that, based on some provable properties, the input graph can be preprocessed so as to detect such motifs and compact them into their corresponding community structures (as dictated by lemmas). This idea would reduce the number of vertices to be processed, which has a direct impact on the overall work of the *biLouvain* algorithm.

In this paper, we prove three such properties: i) for the simple star case (Fig. 12(a); Appendix A.1), ii) for the simple chain case (Fig. 12(b); Appendix A.2), and iii) for the common co-cluster mate case (Fig. 6).

**Fig. 6.**
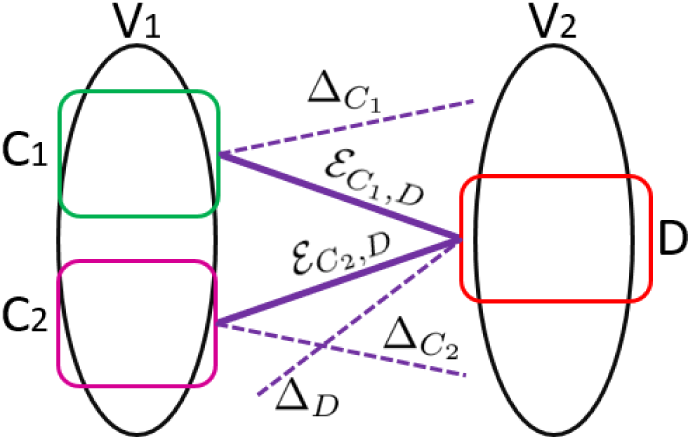
Illustration of the common co-cluster mate case where *ψ*(*C*_1_) = *ψ*(*C*_2_) = *D*.

Although all these properties can be explored conjointly (i.e., they complement one another), we found their effects on performance to be varying. More specifically, based on our tests, we found that identifying simple stars and chains reduces the number of input vertices only by a factor of about 8%. While such an improvement is not necessarily insignificant in itself, relatively, our experimentations with the common co-cluster mate property showed consistently much larger improvements—in some cases, by even more than three orders of magnitude, as will be discussed in Section 6.3.2. Consequently, we delve into the details of the common co-cluster mate property in what follows; and defer the proofs and discussions corresponding to the other two properties (star and chain) to Appendix A.

##### The *Fuse* heuristic

Let *C*_1_ and *C*_2_ be two communities from vertex set *V*_1_, and *D* be a community from vertex set *V*_2_, such that *D* is a co-cluster mate for both *C*_1_ and *C*_2_ (i.e., *ψ*(*C*_1_) = *D*, *ψ*(*C*_2_) = *D*), and (without loss of generality) let *C*_2_ be a co-cluster mate for *D* (i.e., *ψ*(*D*) = *C*_2_). Then the following two lemmas hold:

###### Lemma 5.3.

Given *C*_1_, *C*_2_ and *D* as defined above, the community formed by “fusing” *C*_1_ and *C*_2_ (i.e., *C*_1_ ∪ *C*_2_) will also choose *D* as its co-cluster mate.

***Proof:*** Let:

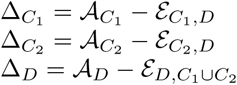

When individual communities *C*_1_ and *C*_2_ have both chosen *D* as their co-cluster mate, their contribution 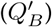 to the overall modularity (*Q*_*B*_) is given by:

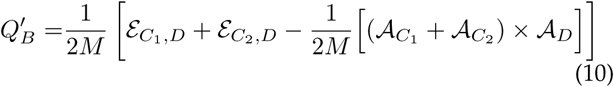

When communities *C*_1_ and *C*_2_ are “fused” together (*C*_1_ ∪ *C*_2_), the contribution of the newly fused community to the overall modularity remains the same as in Eqn. 10. □

###### Lemma 5.4.

Given *C*_1_, *C*_2_ and *D* as defined above, community *D*’s contribution to the overall modularity could potentially increase if *C*_1_ and *C*_2_ are fused (i.e., *C*_1_ ∪ *C*_2_), provided that the condition 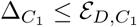 is also met.

***Proof:*** When *C*_1_ and *C*_2_ are individual communities, and *D* has chosen *C*_2_ as its co-cluster mate, then *D*’s contribution 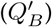 to the overall modularity (*Q*_*B*_) is given by:

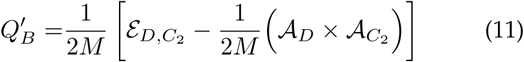

However, if communities *C*_1_ and *C*_2_ are “fused” together, *D* could potentially choose *C*_1_ ∪ *C*_2_ as its new co-cluster mate, in which case, *D*’s contribution to the overall modularity will become:

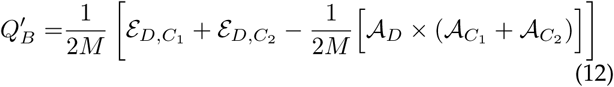

Comparing Eqn. 11 and Eqn. 12:

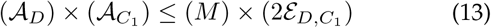

Immediately, from the expression above we notice that:

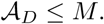

Therefore, community *D* will provide a better modularity contribution choosing *C*_1_ *∪ C*_2_ as its co-cluster mate only if:

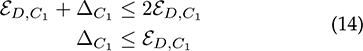

We take advantage of the common co-cluster mate property, shown by the above two lemmas, as follows (see Algorithm 1). By fusing pairs of communities that share a common co-cluster mate, in a preprocessing step, we can possibly reduce the number of initial communities, and in turn, compact the input graph by collapsing the fused communities into vertices. This optimization is aimed at achieving run-time savings; however modularity could potentially be lost if our implementation does *not* explicitly perform the condition in Lemma 5.4. In fact, for the purpose of our implementation, we chose *not* to check for this condition, thereby trading off quality for performance. This makes our implementation of the *Fuse*-based algorithm a heuristic.

Also note that there are multiple minor variations possible while implementing the *Fuse* preprocessing heuristic itself. For instance, a conservative approach is to recompute co-cluster affiliations for communities taken from *V*_2_, after every fuse operation that merges two communities in *V*_1_. However, such an approach is likely to increase the runtime. In our default implementation, we choose a more aggressive approach of recomputing co-cluster affiliations for communities in *V*_2_ only after all fuse operations are completed for communities in *V*_1_. We observed this approach to achieve run-time savings with minimal quality reduction (as shown in Section 6).

##### Run-time complexity for the *Fuse* heuristic

Given that the initial number of communities in a partition is equal to the number of vertices in its corresponding vertex set, the worst-case run-time complexity for assigning co-cluster mates for all communities is 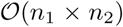, assuming full connectivity as explained in Section 5.3. Then, communities belonging to the same partition are tested for common cocluster mates, for which the worst-case run-time complexity is given by 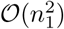 and 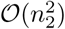, respectively. The final complexity is then 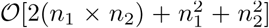 which makes our *Fuse* heuristic a quadratic algorithm. In practice, however, the number of communities drastically reduces (by orders of magnitude as shown in Section 6), thereby making the *Fuse* heuristic highly effective and fast in practice.

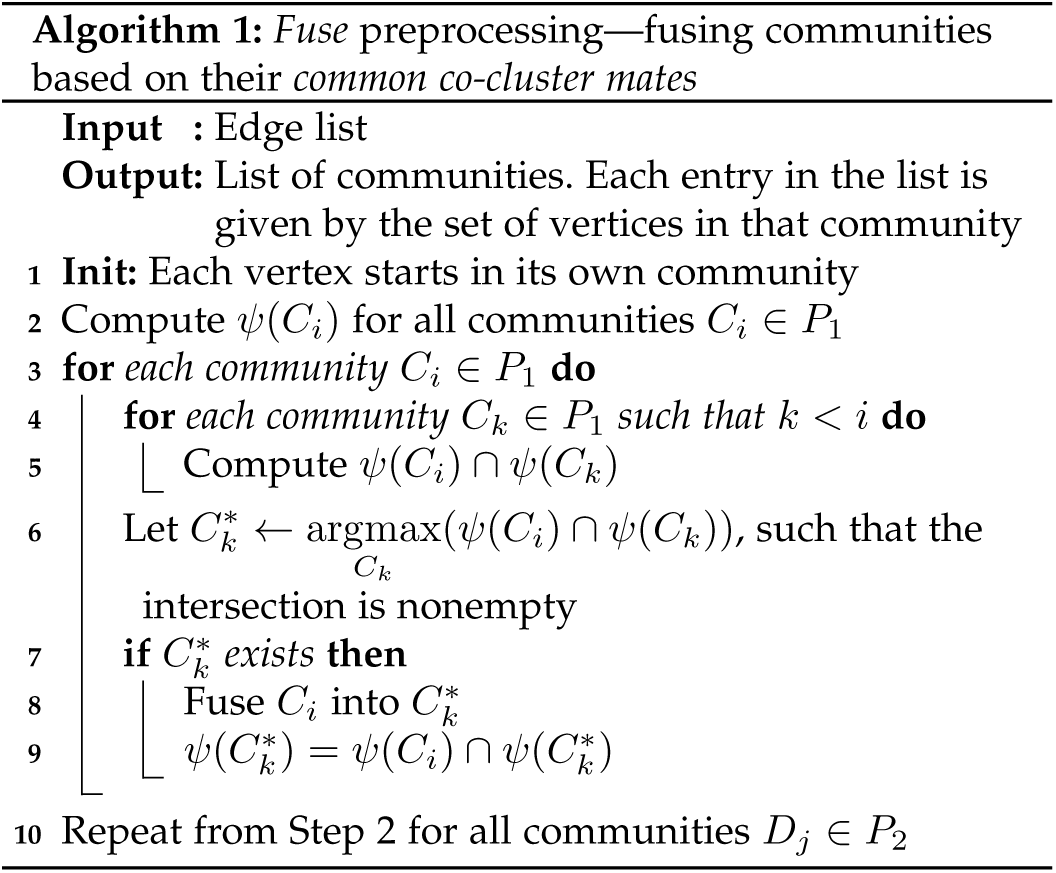

### 5.5 Implementation and Software Availability

We have implemented all steps of our *biLouvain* algorithm (including preprocessing) in C++. Scripts for data format conversions and wrangling were written in Perl. Software is available as open source at https://github.com/paolapesantez/biLouvain.

## 6 Experimental Results

### 6.1 Experimental Setup

#### Experimental Platform

As our experimental platform, we used compute nodes of the *Edison* supercomputer at the National Energy Research Scientific Computing Center (NERSC). Each node has a 12-core Intel “Ivy Bridge” processor at 2.4 GHz, and 64GB RAM. Since our current implementation is serial, it used only one core of a compute node. For graph visualization, we used the Analysis and Visualization of Social Networks (*Visone*) software [23].

#### Test inputs

For experimentation, we used a combination of real world and synthetic data sets (see Table 2):

**TABLE 2.** Bipartite network input statistics.

| Input<br>Data Set | Nodes | | Edges<br>$m$ | Input<br>Type |
| --- | --- | --- | --- | --- |
| | $n_1$ | $n_2$ | | |
| SouthernWomen | 18 | 14 | 89 | Binary |
| memmott1999 | 25 | 79 | 299 | Weighted |
| kevan1970 | 30 | 114 | 312 | Weighted |
| junker2013 | 56 | 257 | 572 | Weighted |
| kato1990 | 91 | 679 | 1,206 | Weighted |
| Malaria | 297 | 806 | 2,965 | Binary |
| Drug-Complex | 680 | 739 | 3,690 | Binary |
| Mouse E135L3 | 163 | 5,162 | 8,870 | Weighted |
| Mouse E115L3 | 159 | 4,341 | 9,770 | Weighted |
| Mouse E155L3 | 156 | 5,301 | 11,237 | Weighted |
| Mouse P28L3 | 89 | 5,924 | 11,253 | Weighted |
| Mouse E185L3 | 146 | 5,339 | 26,735 | Weighted |
| Mouse P4L3 | 128 | 7,796 | 33,001 | Weighted |
| Mouse P14L3 | 230 | 15,302 | 48,242 | Weighted |
| Host-Pathogen | 8,905 | 6,314 | 22,512 | Weighted |
| Gene-Drug | 3,090 | 14,311 | 29,389 | Binary |
| Synthetic1 | 21 | 180 | 216 | Binary |
| Synthetic2 | 67 | 10 | 424 | Binary |

a) *Southern Women [24]:* a women vs. social events bipartite graph;^2^
b) *Plant-Pollinator [5]:* 4 of the 23 pollinator networks, binary and weighted, where an edge represents the frequency of a pollinator’s visit to a plant^3^;
c) *Malaria [22]:* a mapping between subsequences and genes in the malaria parasite *P. falciparum*;
d) *Drug-Complexes [25]:* drug-protein target interactions;
e) *Gene-Drug [4]:* gene-drug interactions;
f) *Genes-Voxels [2]:* mouse brain development during different phases, where edges represent the activation of a gene on a particular point (“voxel”) in three-dimensional space; and
g) *Host-Pathogen [6]:* species-species interactions, where an edge indicates that a pathogen was found in host.

We also used two synthetic networks (*Synthetic1* and *Synthetic2*). More specifically, *Synthetic1* corresponds to a bipartite network with a predefined, well-characterized community structure (probability of 0.9 for intra-cocluster edges and probability of 0.1 for inter-cocluster edges), whereas *Synthetic2* represents a random bipartite network with uniform degree distribution with an unknown (possibly, weak) community structure.

#### biLouvain configuration

All modularity results presented use the *Murata+* formulation defined in this paper (Section 4). Recall that *b*iLouvain has two parameters—the iteration and phase cutoffs (*τ*_*i*_, *τ*_*p*_ respectively), as described in Section 5. We experimented with multiple values of *τ*_*i*_ in the interval [10^*−*6^, 10^*−*2^] on different inputs. These preliminary experiments consistently showed that: a) the final output modularities hardly changed within the interval tested; whereas b) as *τ*_*i*_ is decreased, the number of iterations per phase increased, thereby increasing run-time to completion. Thus, we set the default value of *τ*_*i*_ = 10^*−*2^ throughout our experiments. We set the phase cutoff *τ*_*p*_ to 0.0 in all our experiments. Note that this represents a conservative setting where the algorithm is allowed to terminate only when two consecutive phases produce *n*o change in the overall modularity. We also evaluated the quality-time tradeoff among different vertex ordering schemes in Section 6.3.1. Based on this evaluation, we set *Random* ordering as our default ordering scheme.

### 6.2 Qualitative Assessment

#### 6.2.1 Validation

First, we validate our *biLouvain* algorithm using the Southern Women benchmark and the two synthetic networks. For the Southern Women, our algorithm was able to reproduce the expected communities [26] *identically* as shown in Fig. 7.

**Fig. 7.**
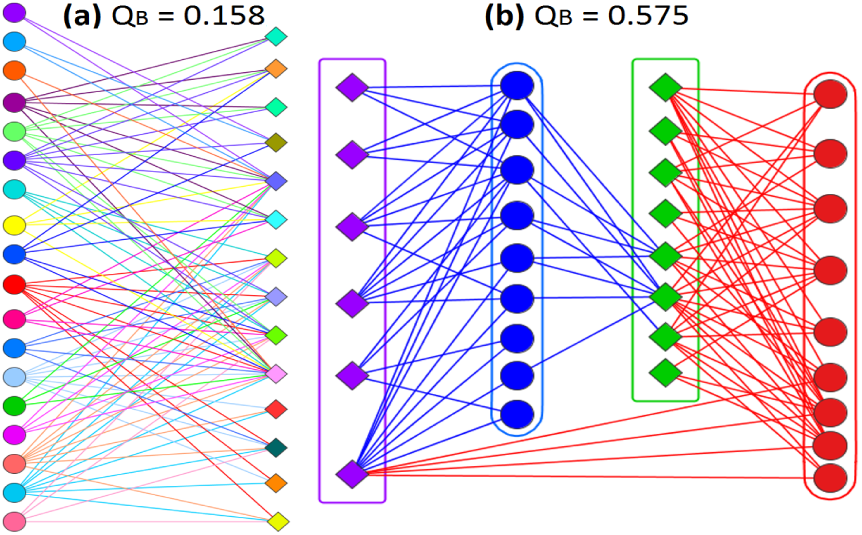
(a) Southern Women Bipartite Graph. Circles - women and diamonds events - by dates. (b) Detected Communities: Red={Evelyn, Laura, Theresa, Brenda, Charlotte, Frances, Eleanor, Pearl, Ruth}, Blue={Verne, Myra, Katherine, Sylvia, Nora, Helen, Dorothy, Olivia, Flora}, Green={6/27, 3/2, 4/12, 9/26, 2/25, 5/19, 3/15, 9/16}, and Purple={4/8, 6/10, 2/23, 4/7, 11/21, 8/3}.

For both synthetic networks, the results were along expected lines. In Fig. 8, we show the *Synthetic2* input, and the bipartite community division output from *biLouvain*. Recall that this is a random network with uniform degree distribution. Yet, our algorithm was able to achieve a modularity of 0.505. On the *Synthetic1* input, which was configured to have a stronger community structure, the output modularity was 0.816 and the expected community structure was successfully recovered.

**Fig. 8.**
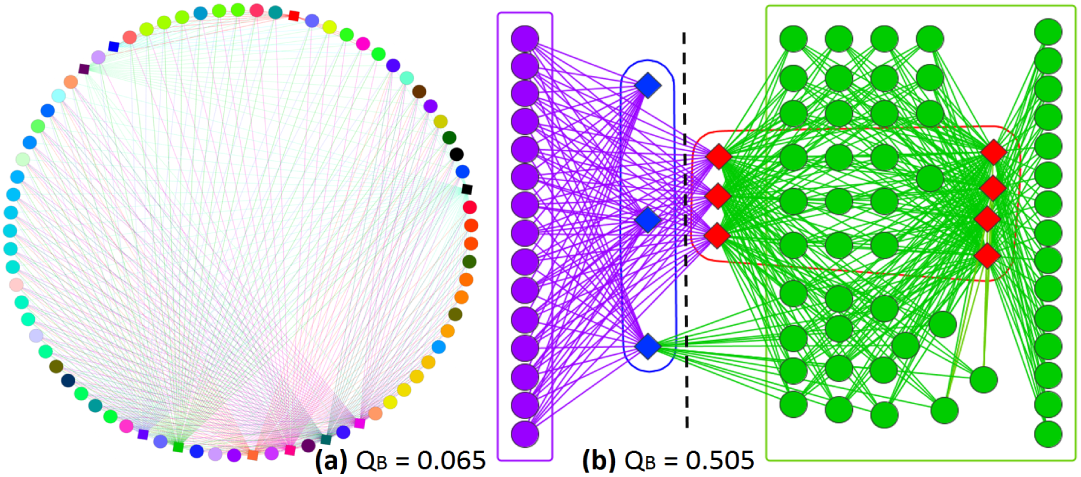
(a) *Synthetic2* bipartite network: Circles represent *V*_1_ vertices, and diamonds *V*_2_ vertices. (b) Bipartite communities output by *biLouvain*: Purple and green vertices form two communities in *V*_1_, while blue and red form two communities in *V*_2_. The dashed line shows the division between the two co-clusters.

#### 6.2.2 Biological Assessment of Clusters

We assessed the significance of the bipartite communities output by the *biLouvain* algorithm, on the Gene-Drug network, which was one of the larger real world networks tested. For assessment, we computed a Gene Ontology (GO)-based significance for each gene cluster detected by our algorithm.

The *biLouvain* algorithm detected 505 gene clusters from the Gene-Drug network, each consisting of two or more genes. We computed the GO significance for these 505 clusters using *gProfileR* [27]. The analysis resulted in 428 (84.75%) clusters with valid GO term annotations. The significance of a particular GO term, associated with a group of genes, is given by its p-value. We used a conservative approach of assigning the *maximum* p-value (i.e., lowest statistical significance) from within each cluster to be the cluster’s p-value. Based on this conservative scheme, we found that *all* of the 428 clusters have a p-value of 0.05 or less—indicating a minimum confidence level of 95%.

The *biLouvain* algorithm detected 505 gene clusters from the Gene-Drug network, each consisting of two or more genes. We computed the GO significance for these 505 clusters using *gProfileR* [27]. The analysis resulted in 428 (84.75%) clusters with valid GO term annotations. The significance of a particular GO term, associated with a group of genes, is given by its p-value. We used a conservative approach of assigning the *maximum* p-value (i.e., lowest statistical significance) from within each cluster to be the cluster’s p-value. Based on this conservative scheme, we found that *all* of the 428 clusters have a p-value of 0.05 or less—indicating a minimum confidence level of 95%.

### 6.3 Performance Evaluation

In this section we evaluate the performance of the *biLouvain* algorithm, including an evaluation of the effectiveness of the different vertex ordering schemes (Section 6.3.1), and of the different heuristics (Section 6.3.2).

#### 6.3.1 Performance of vertex ordering schemes

In Section 5.2 we described three vertex ordering schemes and their potential impact on *biLouvain*’s quality and performance. We studied this quality-time tradeoff on two significantly different real world networks: the Drug-Complex network, which represents the case of an even size distribution between the two vertex partitions (i.e., *n*_1_ ≃ *n*_2_); and the Gene-Drug network, which represents the case of a skewed size distribution between the partitions (here, *n*_1_ ≪ *n*_2_).

Fig. 9 depicts this quality-time tradeoff for these two cases. For Drug-Complex (chart (a)), we observed that all three schemes behave similarly (both by quality and performance). For the Gene-Drug (chart (b)), we observed that the Random scheme demonstrated the best tradeoff, showing a reduction of 0.05 in modularity relative to Sequential, while improving the performance by a factor of 3.43x. These results confirm the expected efficacies of the ordering schemes. These results provide a guide to the choice of the ordering scheme based on the input.

**Fig. 9.**
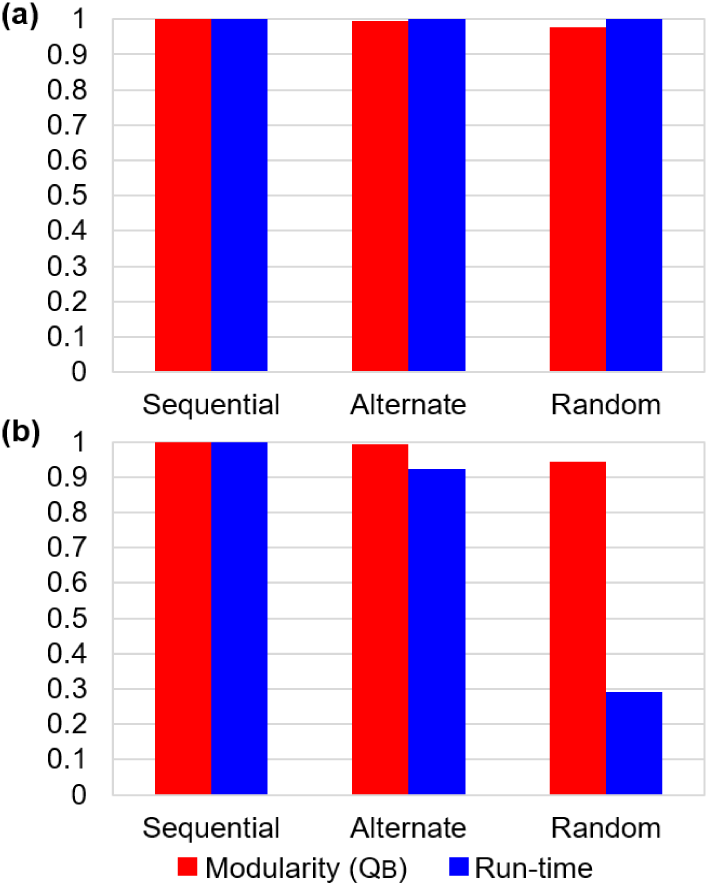
Evaluation of performance of *biLouvain* vertex ordering schemes. (a) Drug-Complex data set (*n*_1_ ≃ *n*_2_). (b) Gene-Drug data set (*n*_1_ ≪ *n*_2_)

#### 6.3.2 Performance evaluation of the heuristics

In this section, we provide a comparative evaluation of two versions of our *biLouvain* algorithm:

- **Baseline** represents the version of our *biLouvain* algorithm that deploys only the random vertex ordering scheme (i.e., without any preprocessing); and
- **Fuse** represents the version which also deploys the *Fuse* preprocessing heuristic that was described in Section 5.4.2.

Table 3 shows the results of comparing the Baseline and Fuse versions of *biLouvain*. Recall that the *Fuse* heuristic essentially is aimed at reducing the number of vertices that are input to the main *biLouvain* clustering algorithm. Consequently we report on both the quality and the runtime performance of both versions. The key observations are as follows.

**TABLE 3.**
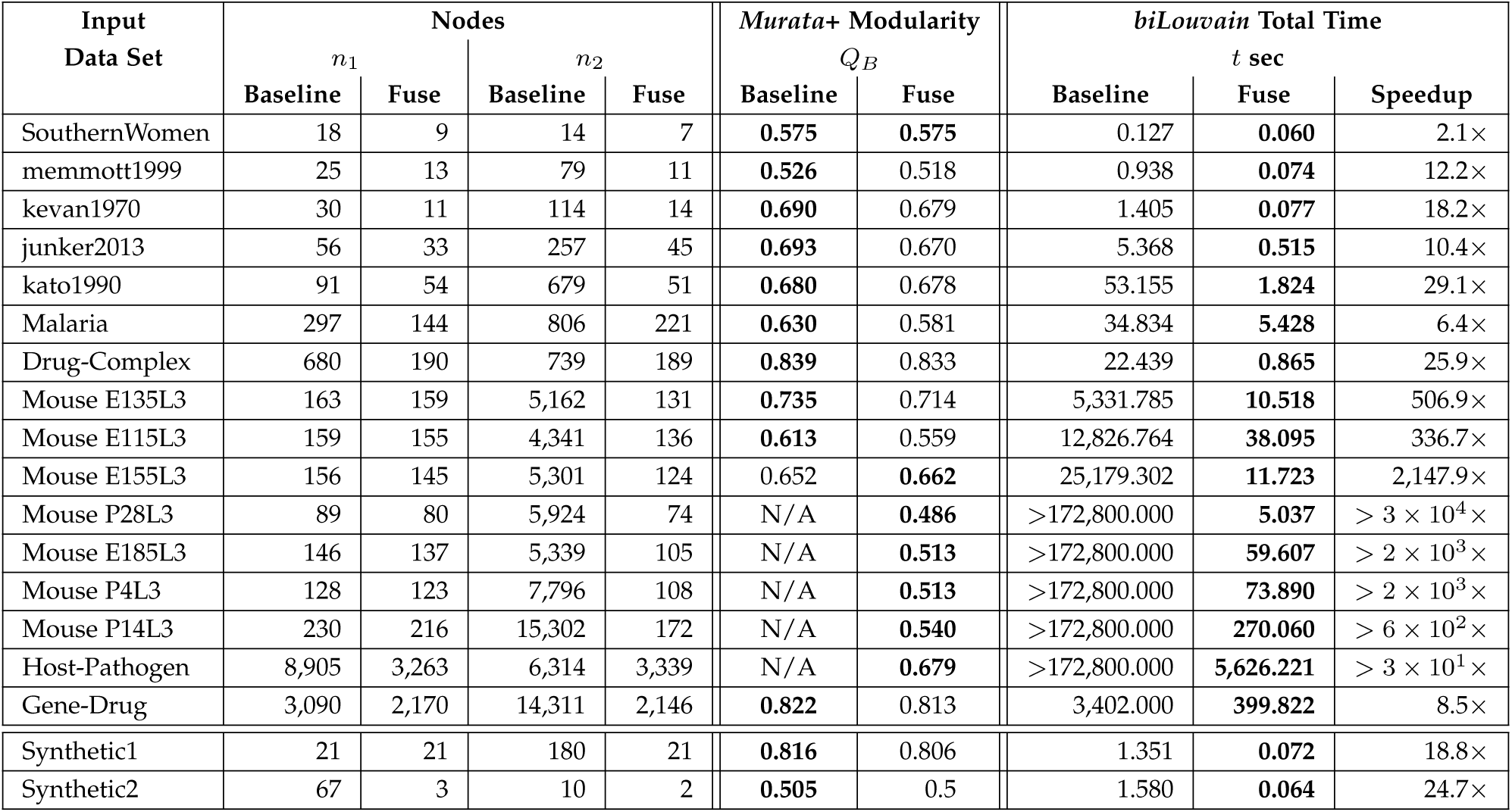
Input statistics, *biLouvain* modularity and run-time performance on the inputs using baseline method and after incorporating the *Fuse* heuristic. The “N/A” label indicates that the results were not available because the corresponding runs did not complete in 48 hours.

The Baseline version takes anywhere between seconds (for the smaller inputs) to hours (for most other inputs), while taking more than 48 hours to completed for 5 out of the 18 inputs tested.

Table 3 also shows the results of running the Fuse version. For all inputs, the Fuse version offers a reduction in the number of vertices—in some cases, up to two orders of magnitude reduction (e.g., Mouse P14L3). This drastic reduction in the graph size meant that we were able to achieve drastic reductions in run-time compared to the baseline version (four orders of magnitude or more in some cases). For instance, the two orders of magnitude reduction in the number of vertices for the Mouse P14L3 meant that the Fuse version was able to complete the computation in less than 5 minutes; compare this with the baseline version not completing in 48 hours. The significant reductions in run-time also resulted in a reduction in the output modularity for many inputs (compared to the Baseline version’s output). However, as Table 3 shows, the loss in modularity is generally marginal. These results show the fine balance in quality vs. time achieved by the Fuse version.

Note that all run-times reported in the Fuse column of Table 3 *include* the time for preprocessing. In fact, we analyzed the cost of *Fuse* preprocessing in Fig. 10. As shown, the cost can be as high as 23% of the total execution time for some input cases; this percentage is a result of the reductions achieved in the subsequent clustering time (compared to the Baseline), thereby demonstrating the high effectiveness of the *Fuse* heuristic.

**Fig. 10.**
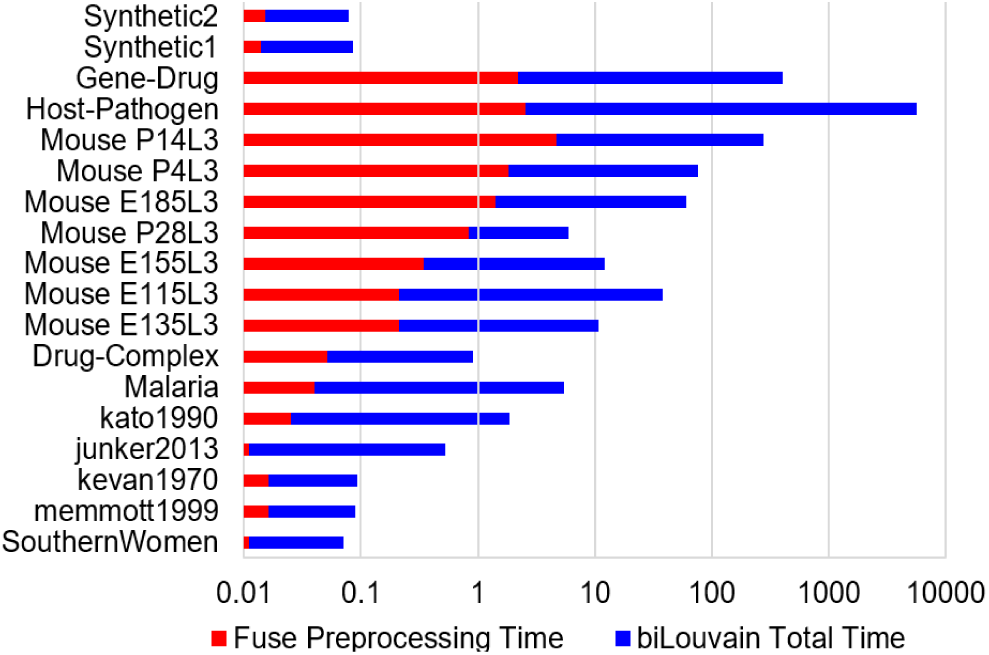
Time taken for the *Fuse* preprocessing step relative to the total time of our *biLouvain* algorithm. Note that the *Fuse* step’s cost is shown as part of the total time, and the time axis is in log scale.

### 6.4 Comparative Evaluation

In this section, we provide a detailed comparative evaluation of *biLouvain* against other existing tools (Section 6.4.1), and a comparison with a projection-based implementation (Section 6.4.2).

#### 6.4.1 Comparison with Other Tools

We compared *biLouvain* against five state-of-the-art methods for bipartite community detection—the Label Propagation (*DIRTLPAwb+*) [20], the Stochastic Block Model (*biSBM*) [22], the *AdaptiveBRIM* [13], the *LPBRIM* [19], and the *Leading Eigenvector* [21]. The latter three algorithms are available in the MATLAB library BiMat [28].

We used the **Fuse** version of our *biLouvain* method in all our comparisons, as it was shown to provide a reasonable quality-time tradeoff in Section 6.3.1.

In our evaluation, we report both on the raw performance (run-time and memory usage) and the quality (based on the Murata+ modularity) of the different methods.

While running *DIRTLPAwb+*, *AdaptiveBRIM*, and *LPBRIM*, we observed that the outputs varied across multiple executions. However, upon a closer examination we found such variations to be minor and therefore we report on an arbitrarily selected output from each of these methods.

In the case of *biSBM*, the outputs not only showed minor variations like the above three methods, but more importantly, the method requires the user to input the number of communities, *k*_1_ and *k*_2_, for the two sides of the bipartite input. In fact, we found that the outputs varied significantly with changes to these values. Therefore, we followed an approach of running the tool over multiple configurations of *k*_1_ and *k*_2_, and selecting the output for which a high modularity was observed while allowing the tool to complete in a reasonable amount of time (i.e., hours). It is to be noted that the run-time of *biSBM* was also highly sensitive to the values of *k*_1_ and *k*_2_.

Table 4 shows the results of our comparative study on individual binary and weighted network inputs. As can be observed, *biLouvain* delivers the maximum modularity for 7 out of the 18 inputs. For the other inputs, the modularity figures delivered by *biLouvain* are generally close to the respective best performing method (except for Malaria). Among the other tools, *Leading Eigenvector* delivers comparable quality to *biLouvain*.

**TABLE 4.** Comparison of *biLouvain* quality and performance against other state-of-the-art methods.

| Input Data Set | Murata+ Modularity $Q_B$ | | | | | | Total Time $t$(sec) | | | | | |
| --- | --- | --- | --- | --- | --- | --- | --- | --- | --- | --- | --- | --- |
|  | Leading Eigenvector | Adaptive BRIM | LP BRIM | DIRT LPAwb+ | biSBM | biLouvain | Leading Eigenvector | Adaptive BRIM | LP BRIM | DIRT LPAwb+ | biSBM | biLouvain |
| SouthernWomen | 0.493 | 0.482 | 0.482 | 0.482 | 0.540 | 0.575 | 1.166 | 0.639 | 0.789 | 0.020 | 0.068 | 0.060 |
| memmott1999 | 0.336 | 0.329 | 0.315 | 0.443 | 0.354 | 0.518 | 0.129 | 0.146 | 0.559 | 3.400 | 0.170 | 0.074 |
| kevan1970 | 0.535 | 0.552 | 0.401 | 0.685 | 0.553 | 0.679 | 0.231 | 0.123 | 1.328 | 5.017 | 0.168 | 0.077 |
| junker2013 | 0.434 | 0.451 | 0.439 | 0.670 | 0.427 | 0.670 | 1.364 | 0.414 | 2.229 | 121.026 | 3.271 | 0.515 |
| kato1990 | 0.513 | 0.476 | 0.485 | 0.655 | 0.467 | 0.678 | 21.674 | 3.123 | 5.131 | 802.217 | 26.977 | 1.824 |
| Malaria | 0.713 | 0.662 | 0.631 | 0.661 | 0.616 | 0.581 | 55.860 | 5.454 | 9.172 | 50,432.690 | 79.268 | 5.428 |
| Drug-Complex | 0.875 | 0.796 | 0.841 | 0.826 | 0.739 | 0.833 | 94.989 | 47.520 | 14.602 | 59,575.560 | 287.388 | 0.865 |
| Mouse E135L3 | 0.772 | 0.685 | 0.678 | N/A | 0.634 | 0.714 | 11,122.779 | 27.701 | 355.864 | >172,800.000 | 3,287.098 | 10.518 |
| Mouse E115L3 | 0.623 | 0.555 | 0.465 | N/A | 0.429 | 0.559 | 4,273.233 | 13.549 | 233.999 | >172,800.000 | 2,023.867 | 38.095 |
| Mouse E155L3 | 0.698 | 0.655 | 0.564 | N/A | 0.524 | 0.662 | 9,417.426 | 19.677 | 369.010 | >172,800.000 | 1,711.001 | 11.723 |
| Mouse P28L3 | 0.539 | 0.480 | 0.481 | N/A | 0.409 | 0.486 | 11,839.985 | 18.197 | 503.175 | >172,800.000 | 2,584.862 | 5.037 |
| Mouse E185L3 | 0.451 | 0.445 | 0.527 | N/A | 0.270 | 0.513 | 10,446.000 | 5.702 | 401.503 | >172,800.000 | 4,170.487 | 59.607 |
| Mouse P4L3 | 0.465 | 0.439 | 0.561 | N/A | 0.284 | 0.513 | 48,607.360 | 13.594 | 1,179.655 | >172,800.000 | 9,911.438 | 73.89 |
| Mouse P14L3 | N/A | 0.412 | 0.412 | N/A | 0.279 | 0.540 | >172,800.000 | 71.866 | 8,446.312 | >172,800.000 | 66,966.088 | 270.060 |
| Host-Pathogen | 0.522 | 0.463 | 0.466 | N/A | 0.390 | 0.679 | 155,031.005 | 15,421.995 | 1,295.048 | >172,800.000 | 35,536.244 | 5,626.221 |
| Gene-Drug | N/A | 0.795 | 0.768 | N/A | 0.825 | 0.813 | >172,800.000 | 3,231.928 | 5,892.775 | >172,800.000 | 23,717.271 | 399.822 |
| Synthetic1 | 0.825 | 0.813 | 0.806 | 0.806 | 0.812 | 0.806 | 0.594 | 0.551 | 1.528 | 17.798 | 1.247 | 0.072 |
| Synthetic2 | 0.453 | 0.456 | 0.454 | 0.456 | 0.456 | 0.500 | 0.090 | 0.033 | 0.731 | 0.032 | 0.069 | 0.064 |

With respect to run-time, Table 4 shows that the *biLouvain* algorithm is the fastest for 10 out of the 18 inputs. More importantly, *biLouvain* consistently delivers one of the fastest run-times for most of the large inputs. Among the other tools, *Adaptive BRIM* delivers a comparable performance to *biLouvain*, although with a significantly lower quality. Furthermore, *Leading Eigenvector*, which delivered a high quality of results in modularity, takes substantially longer run-times on average than *biLouvain*.

To make it easier to see the bigger picture in relative performance (i.e., agnostic to the precise inputs), in Fig. 11 we depict the *performance profiles* of all the methods. These profile charts are constructed as follows: For any input *X*, let the “top” performance (as defined by the desired metric: run-time, quality or memory) was observed for some method *Y*. Then, for each method *Y*’, it’s relative performance on that input is expressed as a percentage of this best performance (by *Y*). For instance, if a method took twice as long time as to complete than the top performing for an input, then the contributing factor 2.0 is added to the list of relative performance values to method *Y’*. Subsequently, the list of all performance values is sorted in non-descending order and plotted on the performance profile chart (one curve for each method).

**Fig. 11.**
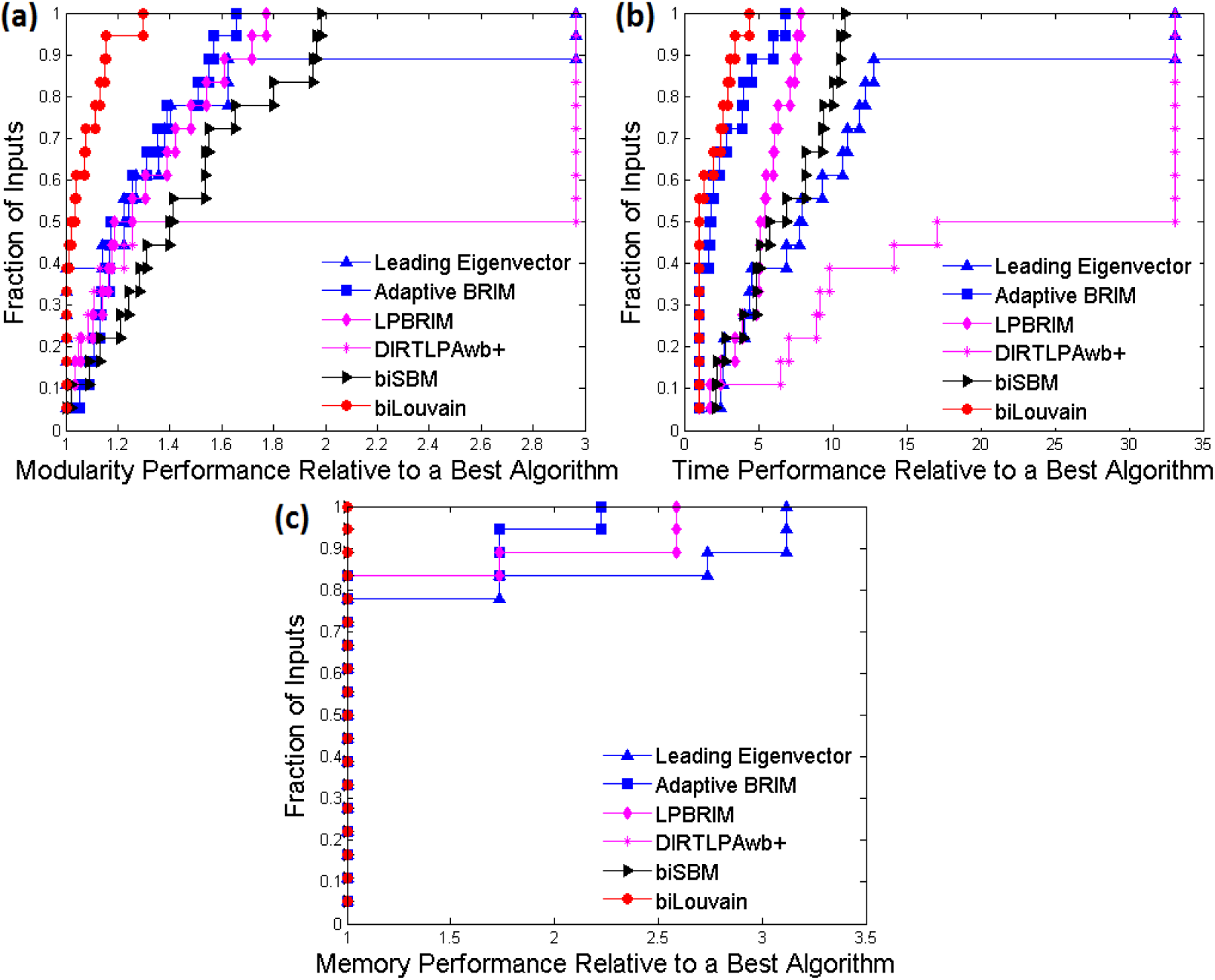
Performance profiles comparing the relative performances of the different state-of-the-art methods versus our *biLouvain* method (Fuse version), over all the 18 input data sets tested. For cases where a method did *not* complete successfully in 48 hours, we assigned a default value either runtime or modularity) that is twice the maximum value observed for that particular input using any other tool. Final modularity scores are shown in chart(a), run-times are shown in chart(b), and memory consumption is shown in chart(c). It is to be noted that the longer a methods curve stays near the Y-axis the more superior its performance is relative to the other methods, over a wider range of inputs.

In the performance plot, the X-axis represents the factor by which a given method fares relative to the best performing method on a particular input. The Y-axis represents the fraction of problems (i.e., inputs). In this scheme of representation, the closer a method’s curve is to the Y-axis, the more superior it’s performance is, relative to the other methods over a wider range of inputs; whereas the worst performance of a method is shown at the top-right most placement of the corresponding curve. Thus, the charts illustrate the relative performance of each method over other methods for the collection of 18 inputs tested (as opposed to the individual inputs).

The performance profile results show that *biLouvain* delivers the highest quality for the most fraction of the inputs (Fig. 11a). Even the worst performance observed by *biLouvain* is less than 1.4x away from the top modularity achieved for that output.

With respect to run-time (Fig. 11b), the performance curve *biLouvain* shows the best results, followed by *Adaptive BRIM*.

With respect to memory consumption (Fig. 11c), *biLouvain* is one of the best performing methods alongside *biSBM* and *DIRTLPAwb+*. We note that, by default, all *biLouvain* runs were performed using a memory limit of 3GB. However, some methods required more memory to complete the execution. For instance, for the Gene-Drug input, *Adaptive BRIM* needed 7GB, *LPBRIM* needed 9GB, and *Leading Eigenvector* required 13GB.

#### 6.4.2 A Projection-based Hybrid Approach

Projection, when applied to bipartite graphs, is known to lose information. However, we explored projection as a potential technique to initialize the set of communities at the start of the *biLouvain* procedure. The goal was to assess the impact of such an initialization procedure on both the quality and run-time of execution.

We implemented our projection-based approach as follows: First, we generate two unipartite graphs—one from *V*_1_ and another from *V*_2_—by simply performing a projection of vertices. Subsequently, for each projected graphs, we run the *Louvain* algorithm [17], which generates a set of communities for each vertex partitions. Using these two sets of communities as “seeds”, we run the *biLouvain* algorithm (Fuse version) on the original bipartite graph inputs. We then compared the output generated by this process against the output generated by running *biLouvain* directly on the input bipartite graph inputs. Table 5 shows the results obtained from using *Projection* in our test input data sets. As expected, we observe an improvement in run-time performance with the use of a projection for community initialization. For instance, we observed two orders of magnitude speedup in the case of the Host-Pathogen input. However, with respect to quality, directly executing *biLouvain* still produces better modularity values for most inputs.

**TABLE 5.**
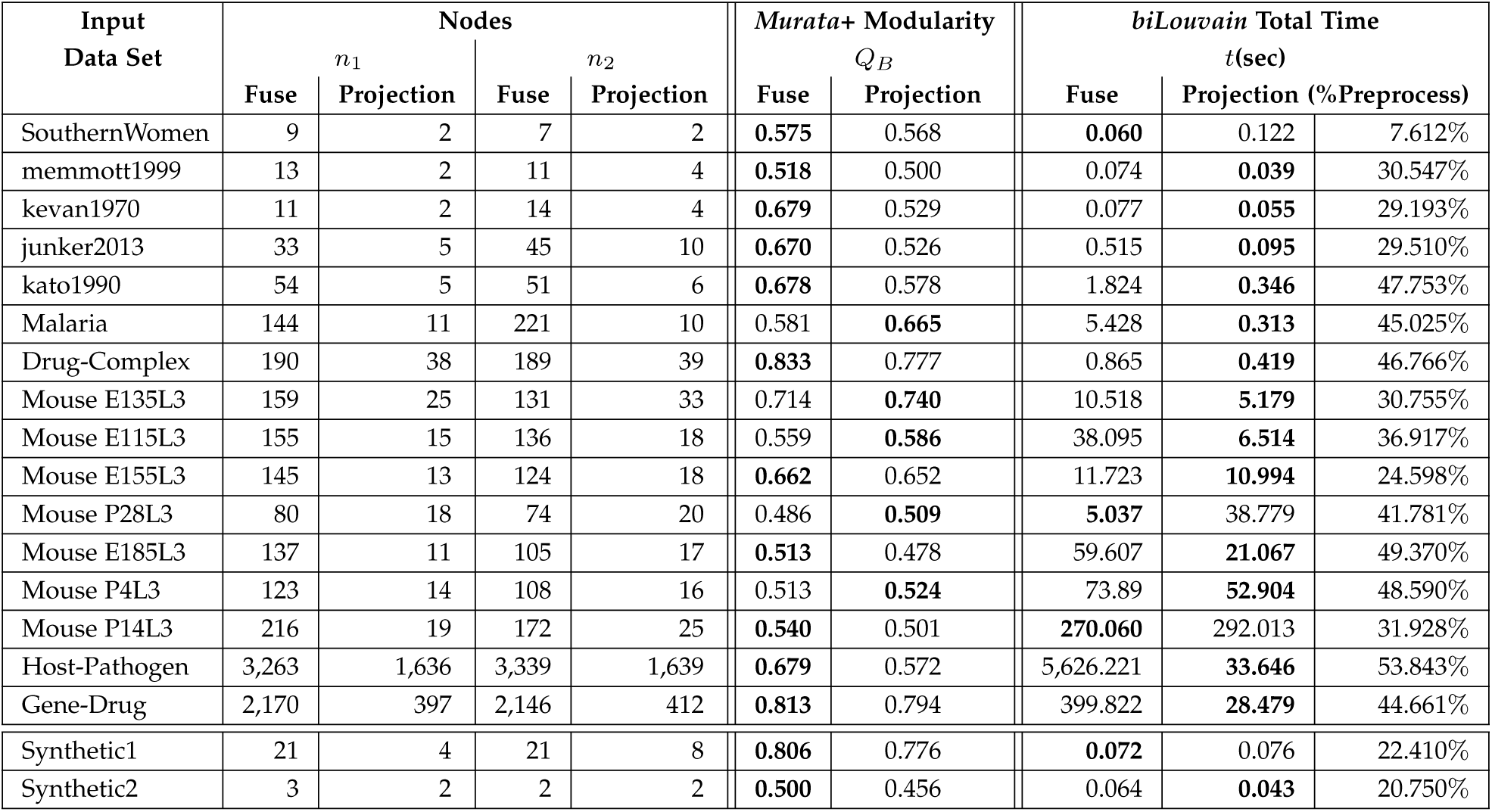
Comparison of our Fuse version of the *biLouvain* to our projection-based hybrid implementation.

Table 6 confirms the expectation that *Projection* all by itself, is not good enough. The table compares the modularities achieved by: (a) if one were to simply derive the communities strictly based on projection (i.e., without running the *biLouvain* step) vs. (b) use the projection-based communities only for initialization and run the *biLouvain* algorithm to compute the final communities. As can be observed, the modularity values produced by the hybrid approach (projection followed by *biLouvain*) is significantly larger than the modularity values produced by projection alone.

**TABLE 6.** Comparison of Murata+ modularities achieved by using Projection-only vs. using the hybrid version that uses projection for community initialization and *biLouvain* for the final communities.

| Input Data Set | Murata+ Modularity |  |
| --- | --- | --- |
|  | Projection-only | Hybrid<br>(Projection-*biLouvain*) |
| SouthernWomen | **0.568** | **0.568** |
| memmott1999 | 0.378 | **0.500** |
| kevan1970 | 0.441 | **0.529** |
| junker2013 | 0.444 | **0.526** |
| kato1990 | 0.431 | **0.578** |
| Malaria | 0.565 | **0.665** |
| Drug-Complex | 0.676 | **0.777** |
| Mouse E135L3 | 0.651 | **0.740** |
| Mouse E115L3 | 0.511 | **0.586** |
| Mouse E155L3 | 0.583 | **0.652** |
| Mouse P28L3 | 0.368 | **0.509** |
| Mouse E185L3 | 0.444 | **0.478** |
| Mouse P4L3 | 0.441 | **0.524** |
| Mouse P14L3 | 0.355 | **0.501** |
| Host-Pathogen | 0.425 | **0.572** |
| Gene-Drug | 0.688 | **0.794** |
| Synthetic1 | 0.612 | **0.776** |
| Synthetic2 | **0.456** | **0.456** |

Furthermore, the projection step in general, requires more memory to complete execution than *biLouvain*—e.g., Mouse P4 (5GB), and Mouse P14 (9GB). This is because the collection of edges leaving a vertex *u* ∈ *V*_1_ (say) in a bipartite graph would create a corresponding clique in the projected subgraph for those vertices in *V*_2_ that it connects to. This may become prohibitive in memory cost for inputs with large vertex degrees.

## 7 Conclusions and Future Work

In this paper, we introduced an efficient algorithm, *biLouvain*, for the problem of community detection in bipartite networks. Our approach is designed to address the dual objectives of minimizing execution time, and maximizing the quality (as measured by bipartite modularity). More specifically, we make the following main contributions: i) (*metric*) We propose a modified variant of the Murata’s bipartite modularity; ii) (*algorithm*) We present a set of efficient heuristics to compute bipartite communities; and iii) (*results*) Our experiments demonstrate the overall runtime effectiveness and qualitative efficacy of the proposed algorithm. The experiments also showed that our algorithm substantially outperforms all the five other existing tools compared in our study, both in execution time (by orders of magnitude) and quality.

Given the paucity in tools for carrying out community detection on bipartite networks, we expect that our method and related software will be of a high utility to the research community. Thus, future extensions have been planned. These include (but are not limited to): i) Parallelization to further reduce the time to solution and enhance problem size reach; ii) Incorporation of intra-type edge information, where available, in addition to inter-type edges as part of modularity computation; and iii) Extending applications on more real world data sets.

## Acknowledgments

This research was supported by US Department of Energy grant DE-SC-0006516. This research used resources of the National Energy Research Scientific Computing Center, a DOE Office of Science User Facility supported by the Office of Science of the U.S. Department of Energy under Contract No. DE-AC02-05CH11231.

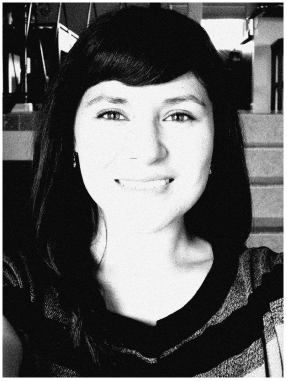

**Paola Pesántez-Cabrera** received the BS in computational systems engineering from University of Cuenca, Cuenca, Ecuador in 2003. She is a 2011 Fulbright scholar. She received her MS degree in computer science from Washington State University, Richland, USA, in 2013. Currently, she is working toward obtaining her PhD degree in computer science at Washington State University, Pullman, USA. Her current research interests include mathematical modeling, algorithm development, computational biology, bioinformatics, graph mining, and data science.

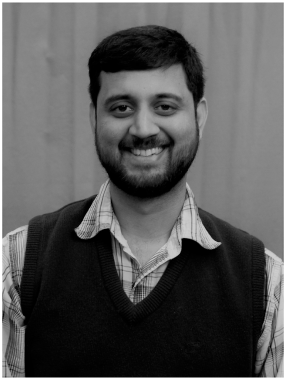

**Ananth Kalyanaraman** received the bachelors degree from Visvesvaraya National Institute of Technology, Nagpur, India, in 1998, and the MS and PhD degrees from Iowa State University, Ames, USA in 2002 and 2006, respectively. Currently, he is an associate professor and Boeing Centennial Chair in Computer Science, at the School of Electrical Engineering and Computer Science in Washington State University, Pullman, USA. His research focuses on developing parallel algorithms and software for data-intensive problems originating in the areas of computational biology and graph-theoretic applications. He is a recipient of a DOE Early Career Award, an Early Career Impact Award and two best paper awards. He serves on editorial boards of IEEE Transactions on Parallel and Distributed Systems and Journal of Parallel and Distributed Computing. Ananth is a member of AAAS, ACM, IEEE, ISCB and SIAM.

## Appendix A Structural Properties

## A.1 Star Property

### Lemma A.1.

A connected component of *G*(*V*_1_ ∪ *V*_2_, *E*, *ω*) that is a vertex *i* ∈ *V*_1_ connected to *k* vertices {*j*_1_, *j*_2_, …, *j*_*k*_} in *V*_2_, is guaranteed to collapse into a single co-cluster.

***Proof:*** Fig. 12(a) illustrates a star input. Let us consider unit weight edges for simplicity. Vertex *i* is guaranteed to be in a community of its own when the algorithm terminates, because all its neighbors (*j*’s) in *V*_2_ are of unit degree. As for the community assignment for all the *j*’s, we consider two possible cases:

*Case a) All the k vertices in *V*_2_ are merged into one community.* For this case, the contribution 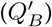 of all these k + 1 vertices, including *i*, to the overall modularity (*Q*_*B*_) is as follows (by Eqn. 5):

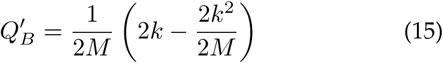

**Fig. 12.**
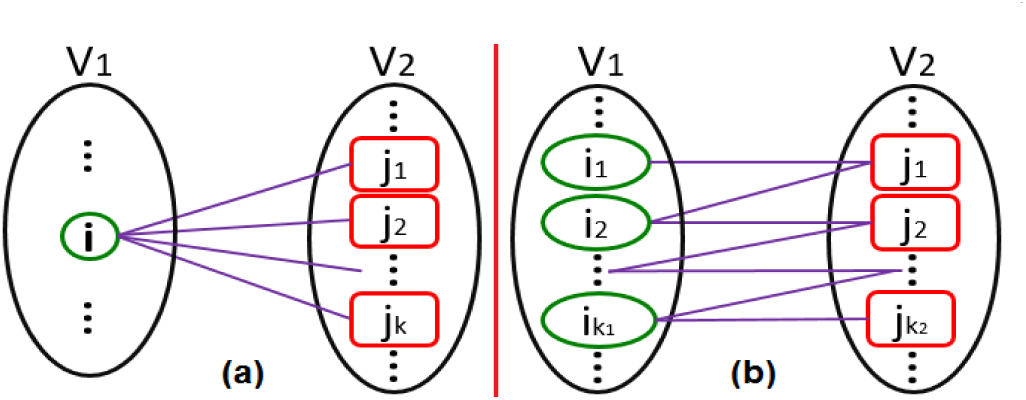
Illustration of a) the star case, and b) the chain case.

*Case b) k* − 1 *vertices in V*_2_ *are merged into one community, the remaining one is left in its own community.* The corresponding modularity contribution is:

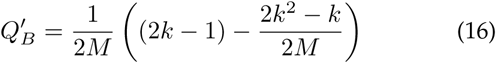

A comparison of Eqn. 15 and Eqn. 16 indicates that the contribution of case (a) can be shown to be always greater than the contribution of case (b), as *M ≥ k* (expression not shown). □

## A.2 Chain Property

### Lemma A.2.

A connected component within *G*(*V*_1_∪*V*_2_, *E*, *ω*) that is a linear chain of length *l* (in the number of edges), can collapse into one co-cluster if and only if 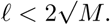

***Proof:*** Fig. 12(b) illustrates a chain input. For convenience we assume unweighted edges. We consider two possible cases to determine the optimal length of a chain for modularity maximization.

*Case a) All vertices forming the chain on each partition, are assigned to the same community.* For this case, the contribution of the chain 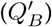 to the overall modularity (*Q*_*B*_) is as follows:

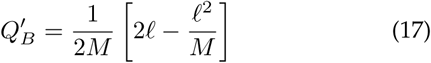

*Case b) The chain is split in two parts, such that community pairs of lengths* 〈*C*_0_, *D*_0_〉 and 〈*C*_1_, *D*_1_〉, *form two co-clusters.* The corresponding modularity contribution is:

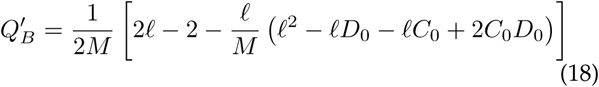

Maximizing the negative term above:

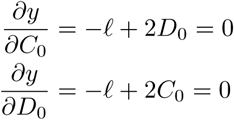

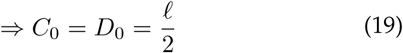

Substituting Eqn. 19 in Eqn. 18:

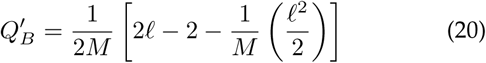

For case (a) to be preferred over case (b):

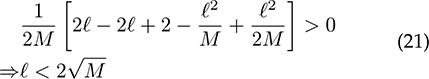

## Footnotes

1. The factor 2 is a result of each edge getting counted twice — once in each direction.

2. This is a bipartite graph with a known community structure and we use this as a benchmark for validation.

3. We experimented on all 23 networks, and select only the top 4 largest networks for presentation in this section.

